# Neural correlates of neonicotinoid-induced impairment of olfactory long-term memory

**DOI:** 10.1101/2022.05.04.489639

**Authors:** Amélie Cabirol, Marco Paoli, Amelie Luise Fehrmann, Nicola Calliari, Marie Fayolle, Sara Bariselli, Jan-Gero Schloetel, Martin Giurfa, Albrecht Haase

## Abstract

Reports of olfactory learning and memory deficits in honey bees exposed to neonicotinoid pesticides have been accumulating over the past decades. As agonists of the nicotinic receptors to acetylcholine, neonicotinoids target most of the projection neurons conveying olfactory information to the mushroom bodies, the brain centres involved in learning and memory. However, the neural mechanisms by which neonicotinoids interfere with memory formation are poorly understood. Here, we investigated the consequences of chronic exposure to the neonicotinoid imidacloprid on the number of projection neuron terminal boutons and on their plasticity in the context of long-term memory formation. Imidacloprid-treated bees exhibited a higher number of boutons in the mushroom bodies, which was associated with long-term memory deficits. Using super-resolution STED microscopy, we characterized synapsin-positive units within boutons and revealed their sensitivity to imidacloprid exposure. Altogether, our experiments suggest that, by altering the number of olfactory projection neuron boutons and their synaptic vesicle content, imidacloprid interferes with the recall of long-term memory.

## INTRODUCTION

Learning and memorizing associations between sensory stimuli and their positive or negative outcome is key to the survival of animals. Associative memory formation, initiated by the coincident detection of a conditioned stimulus (CS) and an unconditioned stimulus (US), triggers the functional and structural reinforcement of synapses located along the neural pathways processing both stimuli ^1–3^. While new synapses are formed between strongly activated neurons, understimulated synapses are weakened and eliminated in a process called synaptic pruning ^4^.

In insects, the mushroom bodies (MBs) are involved in associative odour learning and memory ^5–7^. Within the lip region of the MBs, the terminal boutons of projection neurons coming from the antennal lobes and conveying olfactory information connect with the dendrites of MB intrinsic neurons (Kenyon cells), thereby forming synaptic complexes called microglomeruli (MGs) ^8,9^. Variation in MG number with age and sensory experience has received growing attention ^9,10^. An increase in the number of MGs has been reported in honey bees and fruit flies following paired presentations of an odour (CS) and a sucrose reward (US)^11,12^. In these studies, preventing long-term memory (LTM) formation by unpairing the CS and US stimuli or by inhibiting transcription or protein synthesis precluded such plasticity of projection-neuron axon terminals.

Here we asked whether repeated pharmacological stimulation of projection neurons and Kenyon cells by neonicotinoid pesticides would trigger MG plasticity and interfere with the process of memory formation in honey bees. Neonicotinoids are agonists of the nicotinic acetylcholine receptors ^13^. As such, they target olfactory projection neurons, which are naturally stimulated by acetylcholine in the antennal lobes and also release acetylcholine in the MB lips ^14^. Single doses of neonicotinoids applied directly to neurons *in vitro* or *in vivo* were shown to diminish the response of projection neurons and Kenyon cells to acetylcholine and to odorants ^15,16^. They also decreased the specificity of odour response profiles in the ALs, suggesting that odour discrimination might be affected ^16^. This is consistent with multiple reports of olfactory learning and memory deficits in bees exposed to neonicotinoids ^17,18^. Yet, the causal link between the impact of neonicotinoids on the brain and the observed behavioural alterations still needs to be uncovered by studying both neural and behavioural effects within the same individuals. A recent study showed that a decreased volume of the MB neuropil correlated with poor learning performance in bumblebees following chronic oral exposure to the neonicotinoid imidacloprid ^19^. Still, the volume of a brain region is influenced by multiple parameters (*e.g.*, volume of the dendritic tree and of the axonal arborisations) ^20,21^, and the neuronal mechanisms underlying the impact of chronic exposure to neonicotinoids on the acquisition and storage of olfactory memories are poorly understood ^22,23^.

In the present study, we assessed the impact of chronic oral exposure to imidacloprid on the honey bee MB lip structure and plasticity in the context of olfactory late-LTM formation. Late-LTM is a phase of LTM, observed 72 h after conditioning. It is characterized by its dependence upon *de novo* protein synthesis and structural rearrangements of neurons ^24^. Newborn bees were exposed to one of two concentrations of imidacloprid or to a control sucrose solution for 7 days, before undergoing olfactory conditioning of the proboscis extension response (PER). Combining fluorescent immunostaining of the presynaptic protein synapsin in whole-mount brains with two-photon and STED confocal microscopy, we quantified the number and volume of MGs as well as the density and volume of newly identified synapsin-positive units (SPUs) within MGs. The latter two parameters are relevant proxies of the level of synaptogenesis because synapsin was shown to be abundant in newly formed boutons following neuronal activation in *Drosophila* ^25,26^. Late-LTM formation did not trigger plasticity in the measured neuronal parameters. Yet, reduced LTM performance was observed in imidacloprid-treated bees, which was associated with an elevated number of MGs in the lip. While SPU density was not affected by the imidacloprid treatment, their volume was reduced, indicating that the high number of MGs in the lip of treated bees might not reflect increased synaptogenesis, but rather a deficit in synaptic pruning. Our study, therefore, reveals the importance of maintaining homeostatic MG number in the MB lip for late-LTM in honey bees.

## RESULTS

### Imidacloprid alters the acquisition and long-term storage of olfactory memory

The learning performance of control bees fed a sucrose solution and bees orally treated with imidacloprid (1 ppb or 5 ppb) during their first week of life was assessed using a standardised protocol for olfactory conditioning of the proboscis extension response (PER) (**Fig. 1**) ^27,28^. Bees received five presentations of an odour (conditioned stimulus, CS) and five presentations of a sucrose solution (unconditioned stimulus, US), which were either temporally paired (paired conditioning) or presented independently (unpaired conditioning). The unpaired protocol allowed us to distinguish the effects of repeated CS and US exposure from those specifically attributable to associative learning and memory formation.

**Figure 1.**
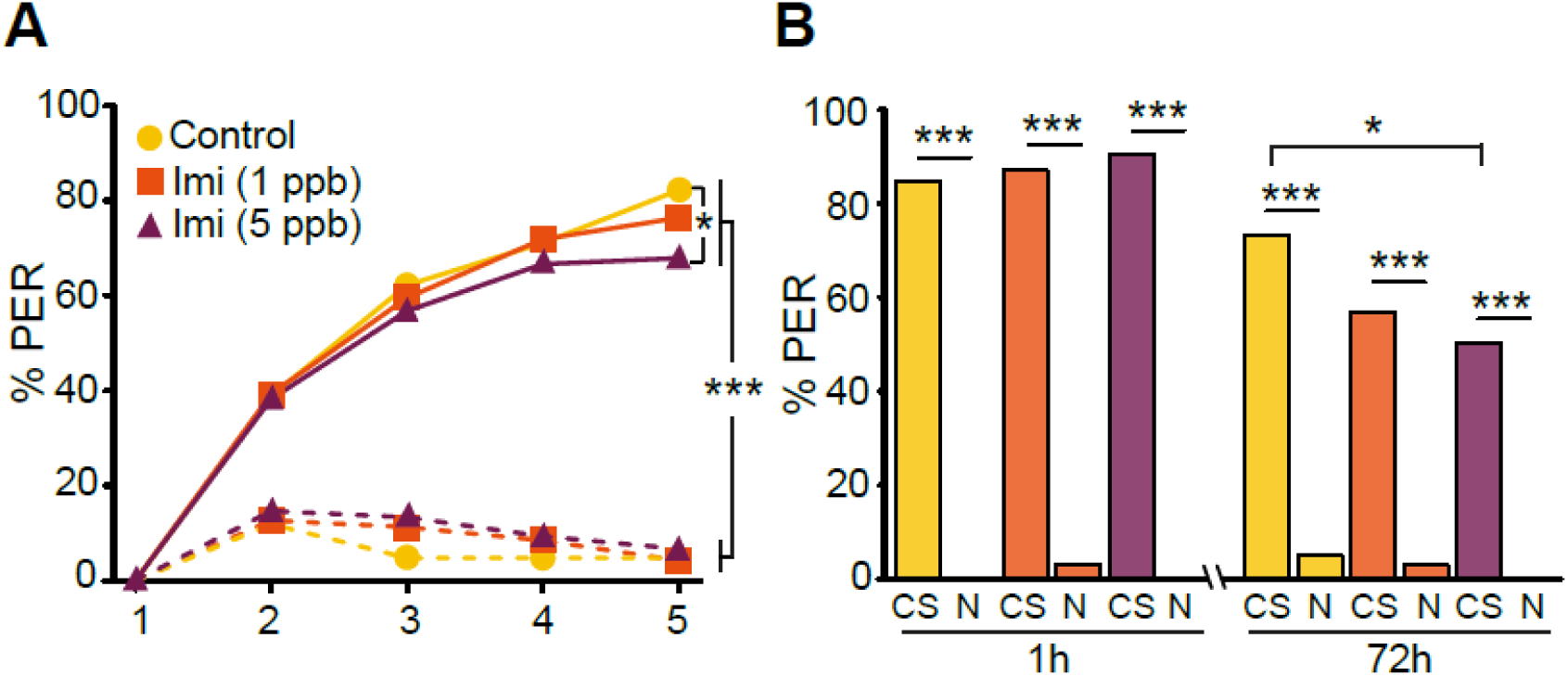
Imidacloprid alters the acquisition and long-term storage of olfactory memory. (**A**) Percentage of bees exhibiting a proboscis extension response (PER) to the conditioned stimulus (CS) across the five learning trials. This percentage progressively increased across trials for the paired groups (solid lines), but not for the unpaired groups (dashed lines). The learning performance of bees exposed to the lowest concentration of imidacloprid (1 ppb; orange) did not differ from the sucrose control group (yellow), while exposure to the highest concentration of imidacloprid (5 ppb; violet) decreased the percentage of bees responding to the CS in the last trial. (**B**) Percentage of learners from the paired group showing PER to the CS and to a new odour (N) during the memory tests performed 1 h or 72 h after conditioning. Bees exposed to the highest concentration of imidacloprid (5 ppb) responded significantly less to the CS than control bees at 72 h after conditioning. (***) *p* < 0.0005, (*) *p* < 0.05; in (A) *n* = 71-89 bees per group, in (B) *n* = 21-36 per group at 1 h and *n* = 34-47 per group at 72 h.

For all treatments, the paired conditioning procedure allowed bees to successfully learn the association between the odour and the sucrose reward (**Fig. 1A**). The percentage of bees extending their proboscis in response to the CS across learning trials differed significantly between the paired and unpaired groups (generalized linear mixed model (GLMM), condition × trial: *χ^2^* = 69.19, *df* = 4, *p* < 10^-4^). In the last learning trial, bees from the paired group responded significantly more to the CS than bees from the unpaired group (GLMM, *post hoc* Wald *z*-tests corrected with the false discovery rate (FDR) procedure; control: *z-*ratio = 11.27, *p* < 10^-4^; imidacloprid 1 ppb: *z-*ratio = 10.31, *p* < 10^-4^; imidacloprid 5 ppb: *z-*ratio = 9.46, *p* < 10^-4^).

The treatment with the highest concentration of imidacloprid (5 ppb) reduced honey bees’ learning performance (**Fig. 1A**). In the last trial, treated bees that underwent paired conditioning responded significantly less to the CS compared to the sucrose control group (GLMM, Wald *z*-tests with FDR correction; *z-*ratio = 2.18, *p* = 0.044). Bees treated with the low concentration of imidacloprid (1 ppb) displayed an intermediate learning performance, which did not differ significantly from either the control (*z-*ratio = 1.18, *p* = 0.32) or the high concentration group (*z-*ratio = 1.04, *p* = 0.38).

To analyse the impact of the imidacloprid treatment on mid-term memory (MTM) and late-LTM as well as its specificity, only bees that solved the learning task during the paired conditioning procedure were successively presented with the CS and a new odour at 1 h (MTM) or 72 h (late-LTM) after conditioning. Bees were not exposed to imidacloprid between the conditioning phase and the memory test, allowing us to assess its impact on memory formation and storage rather than on memory retrieval.

All treatment groups showed specific MTM and late-LTM following the paired conditioning procedure (**Fig. 1B**). Their percentage of PER was significantly higher for the CS than for a new odour (GLM, Wald *z*-tests with FDR correction; *p* < 10^-4^ for all groups). Late-LTM was altered in bees treated with the high concentration of imidacloprid, as shown by their reduced responses to the CS compared to the control (*z-*ratio = 2.33, *p* = 0.03). This was not the case for MTM scores (*z-*ratio = -0.88, *p* = 0.52). Bees treated with the low concentration of imidacloprid did not exhibit significantly different responses to the CS compared to the high concentration group (MTM: *z-*ratio = -0.41, *p* = 0.68, late-LTM: *z-*ratio = 0.77, *p* = 0.44) and to the control (MTM: *z-*ratio = -0.53, *p* = 0.685, late-LTM: *z-*ratio = 1.61, *p* = 0.13). To determine whether the treatment affected generalization of the learned association, we analysed the PER elicited by a novel odorant. Generalization did not differ between groups at either 1 h (Control-Low: *z-*ratio = -0.008, *p* = 1, Control-High: *z-*ratio = 0.00, *p* = 1, Low-High: *z-*ratio = 0.007, *p* = 1) or at 72 h (Control-Low: *z-*ratio = 1.61, *p* = 0.49, Control-High: *z-*ratio = 2.33, *p* = 0.30, Low-High: *z-*ratio = 0.77, *p* = 0.72).

The conditioning procedure had a significant effect on memory retention at 1 h (GLM, condition: *χ^2^* = 137, *df* = 1, *p* < 10^-4^) and 72 h (GLM, condition: *χ*^2^ = 77, *df* = 1, *p* < 10^-4^). Bees from the unpaired group responded significantly less to the CS compared to bees from the paired group (GLM, Wald *z*-tests with FDR correction, *p* < 10^-4^ for all treatments) and did not show significantly different responses between the CS and the new odour at any time point and for any treatment (**Suppl. Fig. S1**). These last results validate the unpaired conditioning procedure and confirm the deleterious impact of the high imidacloprid concentration on long-term memory performance.

### Interactive effects of imidacloprid and memory formation on the lip structure

Next, we assessed whether the impact of 5 ppb imidacloprid on late-LTM was mediated by neuroanatomical alterations in the MB lips. Previous studies have shown that an increase in the number of MGs in the lip was required for late-LTM formation ^12,29^. Following the same protocol, we dissected the brains of bees from the paired group that exhibited specific memory, thereafter called memory performers, and the brains of bees from the unpaired group that showed no PER to either the CS or the novel odour during the 72 h memory test. Importantly, the unpaired group represents a heterogeneous population: because these bees were not subjected to associative conditioning, we cannot determine from their behaviour whether they would have successfully formed LTM had they undergone the paired procedure. Comparing these bees with memory performers therefore reveals neuroanatomical differences associated with successful memory performance. Whole-mount brains were immunostained for synapsin and imaged with a two-photon microscope. The lip volume and the MG density in a subregion of the lip were then measured using an established semi-automated protocol ^30^ (**Fig. 2A-C**). These parameters were used to extrapolate the total number of MGs in the lip.

**Figure 2.**
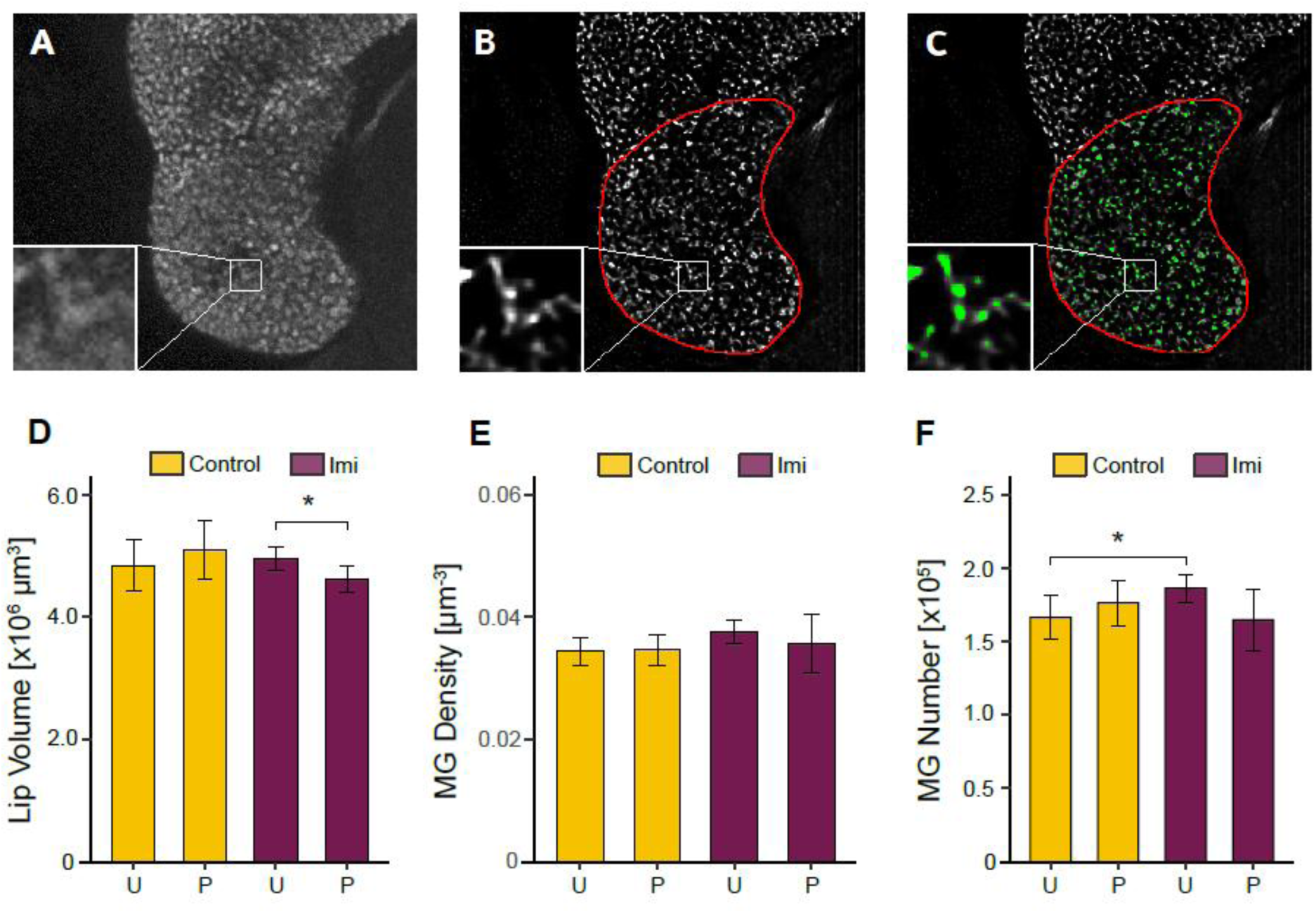
Lip structure is influenced by imidacloprid and long-term memory. **(A)** Two-photon microscopy optical section of a lip immunostained for synapsin. Individual microglomeruli (MGs) can be seen in the inset. **(B)** Image postprocessing (see Materials and Methods section) was applied to improve resolution and contrast. The lip borders (red) were manually delimited on each optical section to reconstruct its volume. The zoomed inset has 10 μm edge size. **(C)** An established segmentation method based on multiple intensity thresholds was applied to detect and automatically quantify the MGs (green)^25^. **(D)** Imidacloprid-treated bees from the paired group (P), which had successfully formed a specific memory of the conditioned stimulus, showed a lower lip volume compared to bees from the unpaired group (U). **(E)** The density of MG in the lip did not differ between groups. **(F)** Imidacloprid increased the total number of MGs in the lip of bees from the unpaired group only. Error bars represent the SEM; (*) *p* < 0.05; *n* = 6-8 per group.

The formation of late-LTM was not associated with structural changes in the lip (**Fig. 2D-F**). Memory performers and bees from the unpaired group did not differ significantly in lip volume (Welch-like ANOVA; *F*(1,24) = 0.95, *p* = 0.34), MG density (*F*(1,24) = 0.03, *p* = 0.87), or MG number (*F*(1,24) = 1.23, *p* = 0.28).

However, significant interactions between conditioning procedure and pesticide treatment were detected for the lip volume (Welch-like ANOVA; *F*(1,24) = 4.27, *p* = 0.0497) and MG number (*F*(1,24) = 6.27, *p* = 0.019). Among treated bees, efficient late-LTM performances were associated with smaller lip volumes. Indeed, memory performers from the paired group had a smaller lip volume than bees from the unpaired group (HC3-robust, Tukey-adjusted pairwise comparisons; *t-*ratio = -2.93, *p* = 0.035), which was not observed in control bees (*t-*ratio = 0.98, *p* = 0.76). It suggests either that a small lip counteracts the effect of imidacloprid on late-LTM, or that the memorization process induced a reduction of the lip volume following imidacloprid exposure.

For MG number, imidacloprid-treated bees exhibited more MGs than control bees, but only under the unpaired condition (*t-*ratio = -2.83, *p* = 0.043). This condition reflects the direct structural effects of imidacloprid on the lip independent of memory formation. No significant effect of imidacloprid was observed in brains collected immediately after the week of treatment (**Suppl. Fig. S2**), suggesting no link between lip structure and the encoding of the CS-US association during conditioning. Instead, the elevated MG number observed after the 72 h memory test in the brain of treated bees seems detrimental to the recall of olfactory memories.

### Imidacloprid alters synapsin-positive units within MGs

Super-resolution STED microscopy allowed analysing the ultrastructure of MGs in the lip of newborn bees and 10-day-old bees, treated or not with 5 ppb imidacloprid during their first week of life (**Fig. 3**, **Suppl**. **Fig. S3**, **Suppl. Video 1-2**). While these 10-day-old bees were not tested in the olfactory conditioning assay, their age corresponds to that of bees having performed the 72 h memory test in previous experiments. The average volume of MGs in subregions of the lip could be measured, as well as the average density and volume of SPUs within the MGs (**Fig. 3A**).

**Figure 3.**
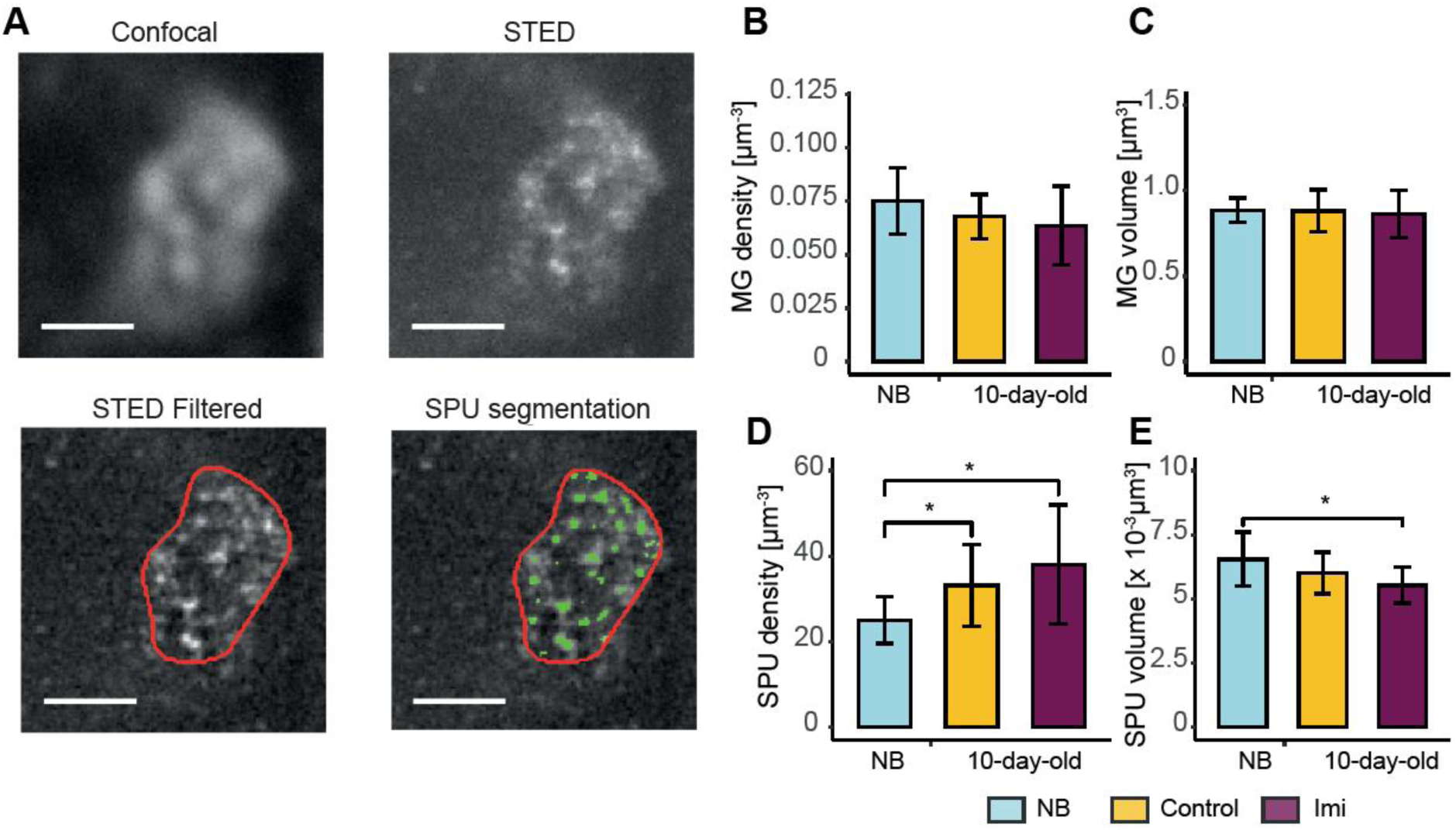
Age- and imidacloprid-related structural changes of microglomeruli. (**A**) Optical section of a MG following α-synapsin immunostaining. Image resolution was improved and MG substructure became accessible via STED. After filtering and delimiting the MG borders (red), the segmentation of synapsin-positive units (SPUs) was performed using an established multiple thresholding algorithm ^25^. Scale bar is 1 µm. (**B**) The density of MGs measured on STED images was not significantly different between newborn (NB; blue) bees and 10-day-old control (yellow) and imidacloprid-treated (violet) bees. (**C**) The volume of MGs was not affected by age and treatment. (**D**) SPU density increased with age. (**E**) SPU volume decreased with age in treated bees. Error bars represent the SEM; (*) *p* < 0.05; *n* = 13-18 per group with 3 experimental cages per group.

While age and imidacloprid treatment did not impact MG density and volume (**Fig. 3B-C**; Welch’s ANOVA; MG density: *F* = 1.54, *p* = 0.24, MG volume: *F* = 0.11, *p* = 0.90), they both affected SPU density (**Fig. 3D**, *F* = 7.10, *p* < 0.005) and volume (**Fig. 3E**, *F* = 3.96, *p* < 0.05). *Post hoc* analyses showed that SPU density was significantly higher in 10-day-old bees compared to newborns, regardless of the imidacloprid treatment (Games-Howell test with FDR correction; control: *p* = 0.016, treated: *p* = 0.030). The volume of SPUs was only reduced in 10-day-old treated bees compared to newborns (*p* = 0.027). Super-resolution analyses thus provided additional information on the effect of age and imidacloprid on MG structure.

## DISCUSSION

Long-lasting stimulation of the nicotinic acetylcholine receptors with imidacloprid increased MG number in the MB lip and reduced olfactory late-LTM performances. Although memory formation was not associated with structural changes in the lip, imidacloprid-treated bees that successfully formed LTM exhibited smaller lips. This may result from an interaction between the plasticity processes induced by LTM formation and those induced by imidacloprid. Alternatively, this brain phenotype may be associated with greater resilience to imidacloprid exposure. Indeed, smaller lips, often reflecting a smaller dendritic tree, may be advantageous to compensate for the imidacloprid-induced high MG number and maintain high LTM capacities. Furthermore, by taking advantage of the increased resolution afforded by STED microscopy, we showed that imidacloprid-treated bees also exhibited smaller SPUs in the lip, thereby providing additional insights into the impact of neonicotinoids on synaptic organization and transmission.

### Imidacloprid induces delayed structural alterations in the mushroom body lip

Consistent with previous reports ^21,31^, the lip volume increased during the first seven days of adulthood. This increase was accompanied by an increase in MG number and likely reflected the ongoing dendritic arborization of Kenyon cells ^32^. While imidacloprid had no immediate effects on the lip structure after the week of treatment, a higher MG number was observed in 10-day-old treated bees exposed to unpaired conditioning. Because the unpaired group allows us to assess the effects of imidacloprid independently of memorization processes, this finding indicates that imidacloprid exposure was associated with a delayed increase in MG number. Notably, this effect was observed three days after the end of treatment. This delayed response suggests that a transient exposure to imidacloprid during a period of active mushroom body maturation interfered with activity-dependent developmental processes whose structural consequences become apparent only later. Such perturbation could also represent a homeostatic response to sustained nicotinic receptor activation, whereby compensatory structural remodelling develops progressively after exposure has ceased^33^.

To determine whether the elevated MG number in the lip of imidacloprid-treated bees reflected synaptogenesis, we quantified the density and volume of SPUs within MGs. Because synapsin is a phosphoprotein that maintains clusters of synaptic vesicles at presynaptic terminals and regulates activity-dependent neurotransmitter release, SPU morphology provides a proxy for presynaptic structural organization ^34,35^. At the larval neuromuscular junction of fruit flies, synapsin was shown to be abundant in new presynaptic boutons and required for their activity-dependent formation ^25,26^. Using STED microscopy, our study revealed an age-dependent increase in SPU density within MGs. This may reflect increased synaptogenesis in the brain of 10-day-old bees compared to newborns. Although SPU density did not differ between imidacloprid-treated and untreated bees, SPU volume was reduced in treated bees compared to newborns, indicating that synaptic vesicle clusters were smaller. Imidacloprid was shown to evoke a concentration-dependent depolarization of Kenyon cells *in vitro,* with detectable effects from 10 ppb onwards ^15^. At the lower concentration used here (5 ppb), the depolarization of neurons might have been too weak and diffuse to enhance activity-dependent synaptogenesis ^2^. Instead, the delayed increase of MG number, combined with smaller SPUs, may reflect deficits in pruning of existing synaptic complexes due to a prolonged and weak stimulation of projection neurons.

### Imidacloprid-associated alterations to MG number may contribute to memory impairment

Chronic exposure to imidacloprid was associated with reduced learning and late-LTM performance, consistent with previous evidence showing that exposure to a field-realistic concentration of 5 ppb impairs olfactory learning and memory in honey bees ^18^. The decreased learning performance in treated bees might have been caused by a reduced sensitivity of projection neurons and Kenyon cells to the release of acetylcholine triggered by the olfactory stimulation ^15,16^. A deficit in synaptic pruning in the lip of treated bees, resulting in a high MG number, is a plausible cause for their reduced late-LTM performance. While in the unpaired group, imidacloprid-treated bees had a higher MG number than control bees, this was not the case in memory performers from the paired group. Long-term memory formation relies on the creation of new synaptic connections between the most activated neurons of the network and on the pruning of the least activated connections ^2^. The non-specific activation of projection neurons by imidacloprid during the conditioning may have prevented the synaptic pruning required for the consolidation of the learned CS-US association. Another possibility is that the high number of MGs in treated bees interfered with the retrieval of the learned CS-US association during the memory test. Previous studies have shown that a high number of MGs in the lip was associated with poor performance in an olfactory reversal learning task ^31,36^. This might be explained by a decreased sparseness of olfactory responses in the MBs. Sparse coding of olfactory information is an important property of the insect MBs ^37–40^. The firing of Kenyon cells is highly odour-specific and allows the discrimination of similar odours in fruit flies ^41^. Olfactory long-term memory formation was shown to increase both the number of MGs on the projection neurons encoding the CS ^42^ and the number of Kenyon cells responding to the CS ^43^. The elevated number of MGs in the lip of imidacloprid-treated bees in our study might have led to the activation of a larger number of Kenyon cells by the CS during the memory test, thereby reducing stimulus specificity and impairing retrieval of the learned CS–US association.

### Long-term memory formation was not associated with changes in lip structure

Previous work showed that MG density in the mushroom body lip increased 72 h after olfactory conditioning using the PER paradigm ^11^. Surprisingly, we did not detect comparable changes despite following the same conditioning protocol and experimental design as Hourcade *et al.* (2010)^29^. One methodological explanation is that we used a different approach to quantify MG density, sampling a larger subregion of the lip ^44^. In *Drosophila*, an increase in MG number was shown to be specific to the projection neurons activated by the CS ^42^. Since odours were shown to specifically activate a sparse number of MGs in honey bees ^40^, likely, the structural changes occurring in a subset of projection neurons were not detectable when quantifying the overall MG density. In other words, the few 2D sampling regions used by Hourcade *et al*. (2010)^29^ may have coincidentally included a higher proportion of MGs undergoing structural changes, whereas averaging across the larger volume sampled here may have diluted such localized effects.

### A small lip is associated with good olfactory LTM performance under imidacloprid treatment

In addition to the alterations in MG number and SPU morphology, we found that imidacloprid-treated bees that successfully formed long-term memories had smaller lips. This finding is particularly interesting in light of previous evidence linking mushroom body volume to cognitive performance. In bumblebees, the same concentration of imidacloprid reduced olfactory learning performance and MB volume after only 3 days of chronic exposure ^19^. The authors found a positive relationship between MB volume and learning performance. In honey bees, MB volume was also found to be positively correlated with cognitive performance ^45^. Our results point in the opposite direction, with memory performers having smaller lips than unpaired bees, a heterogeneous group comprising individuals with potentially different memory abilities. This may be due to a different treatment duration, a different timing for brain dissections, or to our focus on the olfactory neuropil of the MBs. Additionally, the relationship between brain volume and cognitive performance is unlikely to be straightforward. Volumetric variation can arise from different neuronal processes, including dendritic arborization and synaptic pruning ^20,21^.

Two explanations may account for our findings. First, smaller lips may reflect structural remodelling associated with successful long-term memory formation under imidacloprid exposure. Memory-related plasticity may interact with the effects of imidacloprid on neuronal organization, resulting in a distinct structural phenotype among treated memory performers. Second, smaller lips may represent a pre-existing anatomical feature associated with greater resilience to imidacloprid. For example, a smaller dendritic tree might reduce the consequences of the increased MG number associated with exposure, potentially helping preserve the selectivity of olfactory representations.

### Conclusions

Our study combined cognitive assays and super-resolution microscopy to uncover the neural mechanisms by which neonicotinoid pesticides affect olfactory memory in honey bees. We found that sustained activation of nicotinic receptors resulted in a high number of MGs in the lip, which was detrimental to LTM performance. Future studies should determine whether this increase in MG number resulted from a disruption of activity-dependent synaptic pruning of cholinergic neurons induced by the pesticide.

## MATERIALS AND METHODS

### Animals

Experiments were performed from May to October on honey bees (*Apis mellifera*) collected from one hive maintained at the University of Trento in Rovereto (Italy). Newborn adult bees were obtained by placing a comb of sealed brood in an incubator (34°C, 55% humidity) overnight. On the following morning, bees that had emerged from their cell were placed in cages (5 × 8 × 5 cm^3^; 15 bees per cage) and left in darkness in an the incubator for 7 days. Honey bees for whole-brain labelling and subsequent STED analysis were collected from colonies maintained on the roof of Sorbonne University, Jussieu Campus, Paris (France).

### Imidacloprid treatment

In the cages, bees had unlimited access to water and sucrose solution (50% w/w). In some cages, the sucrose solution was supplemented with imidacloprid (1 ng/g or 5 ng/g; Sigma-Aldrich). A stock solution of imidacloprid (100 ng/g) was prepared and stored at -20°C. The water and treatment solutions were changed every day.

### Olfactory conditioning

At the end of the 7th day of treatment, the sucrose solution (± imidacloprid) was removed from the cages. Two hours later, bees were immobilized on ice and harnessed in small metal tubes allowing movement of the antennae and mouthparts only. They were kept in a dark, humid chamber at room temperature overnight.

The olfactory conditioning experiment started on the following morning and followed the same protocol as in Hourcade *et al.*^46^. Bees that did not extend their proboscis in response to a 50% sucrose solution (w/w) were discarded prior to the conditioning experiment. Individual bees were placed in front of an olfactometer delivering a continuous airflow for 15 s (familiarization with the context). The conditioned odour (1-hexanol, >99 % pure, in a 20 mL glass vial; Sigma-Aldrich) was then delivered for 4 s and the sucrose reward for 3 s, with a 1 s overlap between both. The bee was left in front of the airflow until the end of the learning trial. The duration of each trial was 40 s, and the inter-trial interval was 10 min. Bees received 5 presentations of an odorant and 5 presentations of a sucrose reward in combination (*paired group*) or not (*unpaired group*). Bees from the *unpaired group* cannot learn the association between the odour and the sucrose reward, which are not presented in combination, and this group, therefore, represents a control for the effect of repeated exposure to an odorant and a sucrose reward. To balance the amount of time spent on the set-up between both groups, bees from the *paired group* also received five blank trials during which they were placed in front of the airflow for 40 s without any stimulation ^47^ . The order of presentation of reinforced, blank, odorant-only, and sucrose-only trials was randomized.

For bees that showed successful learning, by exhibiting PER to the CS during the 5th trial, memory was assessed either 1 h (mid-term memory, MTM) or 72 h (late long-term memory, late-LTM) after conditioning. Bees tested for MTM were left in darkness, at room temperature, for 1 h after the conditioning experiment. Bees tested for late-LTM were placed back into cages (10 bees per cage) with sucrose solution and water *ad libitum*, in an incubator (34°C, 50% humidity). On the day before the LTM test, bees were harnessed again in metal tubes at 5 p.m. and left in darkness at room temperature overnight. Both memory tests (MTM and late-LTM) consisted of the presentation of the conditioned odorant using the same design as for the conditioning experiment. A new odorant (1-nonanol, >98% pure; Sigma-Aldrich) was also presented in a different trial to assess odour generalization. The order of presentation of the conditioned and the novel odorant was randomized between bees. At the end of the memory test, the integrity of the PER was assessed for each bee by touching the antennae with a toothpick soaked in sucrose solution; bees not responding with PER to this stimulation were discarded from the analysis.

### Immunostaining of whole-mount brains

Immediately after the late-LTM test, the brains of bees from the *paired group* that had formed a memory, and the brains of bees from the *unpaired group* that had not formed memory were dissected in cold saline (130 mM NaCl, 5 mM KCl, 4 mM MgCl_2_, 5 mM CaCl_2_, 15 mM Hepes, 25 mM glucose, 160 mM sucrose; pH 7.2). For the experiment using STED microscopy, the brains of newborn bees and 10-day-old bees treated with 5 ppb imidacloprid during their first week were dissected. The trachea and membranes surrounding the brain were removed, and the brain was fixed overnight in paraformaldehyde (4% in PBS) at 4°C on a shaker. The protocol for staining synapsin in whole-mount brain followed Groh *et al.* ^48^. After fixation, brains were rinsed with PBS (0.01M; pH 7.2; 3×10 min), permeabilized in 2% Triton x-100 (in PBS: PBS-Tx; 1×10 min) and in 0.2% PBS-Tx (2×10 min), and finally blocked in 2% normal goat serum (NGS; in 0.2% PBS-Tx) for 1 h. All these steps were performed at room temperature, on a shaker. Brains were then incubated with the primary antibody directed against synapsin (SYNORF1, DSHB; 1:10 in 0.2% PBS-Tx, 2% NGS) for four days at 4°C, on a shaker. After rinsing the brains in PBS (5×10 min) at room temperature, brains were incubated for three days at 4°C with either the Alexa Fluor 488-conjugated goat anti-mouse secondary antibody (Fisher Scientific, 1:250) for subsequent two-photon imaging, or the Abberior STAR RED-conjugated antibody (1:200) for STED imaging. Brains were rinsed again in PBS (5×10 min) and dehydrated in an ascending ethanol series (30%, 50%, 70%, 90%, 95%, 100%, 100%, 100%; every 10 min). Finally, they were cleared and mounted in methyl salicylate for imaging.

### Two-photon imaging

Images of whole-mounted brains were acquired with a two-photon microscope (Ultima IV, Bruker), equipped with a 20× objective (NA 1.0, water immersion, XLUMPLFLN20XW, Olympus) and a Ti:Sa laser (Mai Tai Deep See HP, Spectra-Physics) tuned to *λ* = 780 nm. For all images, the resolution was 512 × 512 pixels and the pixel dwell time was 30 µs. For volumetric measurements, optical sections of the right median calyx were taken every 5 µm throughout the entire structure with a pixel size of 1.03 × 1.03 µm^2^. For MG quantification, a subregion of the outer lip of the right median calyx was then imaged every 0.5 µm across 50 µm with a pixel size of 0.34 × 0.34 µm^2^. Depth-dependent signal attenuation was compensated by increasing the laser power automatically along an exponential profile.

### STED imaging

Whole-mounted brains were imaged with a stimulated emission depletion (STED) super-resolution microscope (Expert Line, Abberior Instruments) using a 100× objective (NA 1.4, oil immersion, UPLSAPO100XO, Olympus), pulsed excitation at 640 nm and pulsed depletion at 775 nm (**Suppl. Fig. S3**). Constant image quality in shallow as well as deeper sample regions was ensured by compensation of spherical aberrations in a depth-dependent manner via a deformable mirror (Adaptive Optics module, Abberior Instruments). Within the outer lip of the right median calyx, two volumes (XYZ, volume size 10 × 10 × 9.9 µm^3^ and voxel size 25 × 25 × 300 nm^3^) were imaged with a dwell time of 7.5 µs and a pinhole size of 0.7 AU (**Suppl. Video 1**). Confocal and STED images were acquired in line-sequential mode, summing intensity values from 2 repeated line scans (confocal) or 6 line scans (100% 2D-STED PSF). Imaging was performed at the Institut du Cerveau, Hôpital Pitié-Salpêtrière, Paris (France).

### Image processing and analysis

Two-photon image stacks were post-processed by 3D deconvolution with a measured point spread function (PSF) in AMIRA (Thermo-Fisher, v2019), using the Extract Point Spread Function and Deconvolution module. The image stacks were resampled to obtain an isotropic voxel size of 0.1 × 0.1 × 0.1 µm^3^. The total volume of the lip was measured in AMIRA after manual delimitation of the lip borders. A similar delimitation was performed on images of a subregion of the outer lip used for MG quantification. The segmentation of individual MGs within the resulting lip masks was performed using a custom code in MATLAB (Mathworks, v.2021) following a multiple-threshold approach to ensure reproducible detection across different imaging conditions ^30^. Ninety normalized intensity thresholds ranging from 0 to 0.23 were applied to the image stacks to identify three-dimensional regions of connected voxels above each threshold (MG candidates). Only MG candidates with volumes between 3·10^-3^ µm^3^ and 5 µm^3^ were considered for further analyses. Regions detected at different thresholds were considered part of the same MG if their centroids were within 0.4 µm and the region detected at a higher threshold was contained within the corresponding region detected at a lower threshold. The larger MG candidate was retained. This allowed individual MG candidates to be distinguished independently of the choice of a single intensity threshold.

All STED image stacks were post-processed using a difference of Gaussians (DoG) approach. They were drift-corrected prior to further processing. Image stacks were resampled to an isotropic voxel size of 0.025 × 0.025 × 0.025 µm^3^ by three-dimensional interpolation. Segmentation was performed with the previously described multiple-threshold algorithm at 22 intensity thresholds distributed between normalized intensity values of 0.2 and 0.55. MG candidates with volumes ranging between 4.64·10^-5^ µm^3^ and 2 μm^3^ were retained. Candidates were considered to represent the same MG if they were detected at 2 different thresholds and their centroids were within 1.4 µm. When several MG candidates represented the same MG, the MG volume was determined by the largest volume of the candidates. Individual MG masks were generated for further analyses.

The microglomerular ultrastructure within the previously identified STED MG masks was segmented using the multiple-threshold algorithm described for the two-photon analysis. Individual synapsin-positive units (SPUs) were extracted by applying 70 intensity thresholds between 0.05 and 0.8. SPU candidates with volumes between 5.41·10^-4^ µm^3^ and 2·10^-2^ µm^3^ were retained. Detections obtained at different thresholds were considered to represent the same SPU when their centroids were within 0.05 µm, and the region detected at a higher threshold was contained within the corresponding region detected at a lower threshold. The larger SPU candidate was then retained. Additionally, mean volumes of segmented SPUs were determined for each MG. Measurements performed on two subregions of the lip were averaged for each individual.

Parameters and segmentation criteria were optimized separately for MG segmentation of two-photon and STED image stacks and for SPU segmentation, based on a comparison with manual counts obtained in a subset of ten brains. All algorithms were implemented in MATLAB.

### Statistical analyses

The presence or absence of PER to the CS and to the new odour during the olfactory conditioning and memory tests was recorded as a binomial variable. Different generalized linear mixed models with a binomial error structure — logit-link function — (glmer function of R package lme4) were used to assess the predictive value of the trials, conditioning procedure (paired/unpaired), and treatment on individual responses during the acquisition phase. The individual identity was set as a random factor. When necessary, models were optimized with the iterative algorithm BOBYQA ^49^. Generalized linear models with a binomial error structure — logit-link function — were used to assess memory performance. Model selection was performed by retaining the significant model with the lowest Akaike information criterion (AIC) value. When AIC values were very similar, the most significant model was retained. *Post hoc* pairwise comparisons of estimated marginal means were performed using Wald *z*-tests, with *p*-values adjusted for the false discovery rate (FDR).

The normality of residuals and homoscedasticity of the lip, MG, and SPU measurements were assessed with a Shapiro-Wilk and Bartlett test, respectively. ANOVAs – or Welch’s ANOVAs in case of unequal variances – were then applied to assess the impact of the treatment, conditioning procedure, and age on these variables. Post-hoc pairwise comparisons of estimated marginal means were performed using HC3 robust standard errors, with Tukey adjustment for multiple comparisons.

## DATA AND CODE AVAILABILITY

Unprocessed datasets and R codes are available on Github at acabirol/Neonicotinoid-induced-memory-impairment. The MATLAB segmentation analysis code is available on https://github.com/NeurophysicsTrento/BeeMushroomBodySegmentation/.

## ACKNOWLEDGEMENTS

The study was funded by the Autonomous Province of Bolzano (A.H.; project TN2214 STEFANIE), the CNRS (A.C.) and the ERC Advanced Grant Cognibrains (M.G.; project 835032).

## AUTHOR CONTRIBUTIONS

A.C., M.P. and M.F. conducted the experiments; A.C., M.P. and G.S. acquired the STED images; A.C., A.F., N.C. and S.B. conducted the image and statistical analyses; A.C. and A.H. designed the experiment and drafted the manuscript; A.C., A.H. and M.G. provided funds and resources; all authors contributed to finalizing the manuscript.

