## Supplementary information for "Neural correlates of neonicotinoid-induced impairment of olfactory long-term memory"

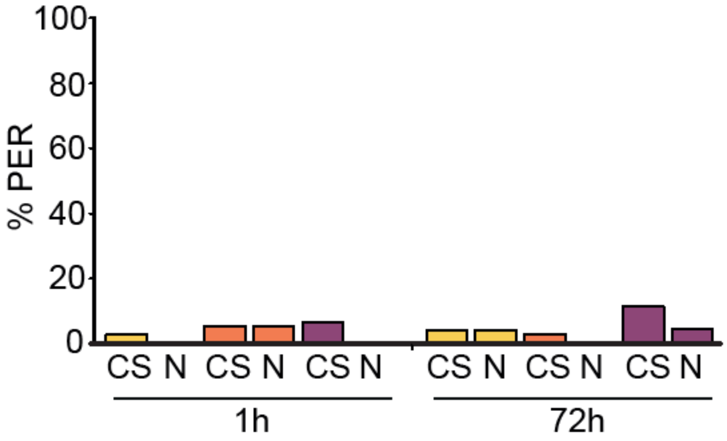


**Supplementary Figure S1. Olfactory memory performance of bees from the unpaired groups chronically exposed to imidacloprid.** The percentage of control bees (yellow) and treated bees (1 ppb, orange; 5 ppb, violet) exhibiting a proboscis extension response (PER) did not differ significantly between the conditioned stimulus (CS) and a new odour (N) at 1 h and 72 h after the conditioning experiment.

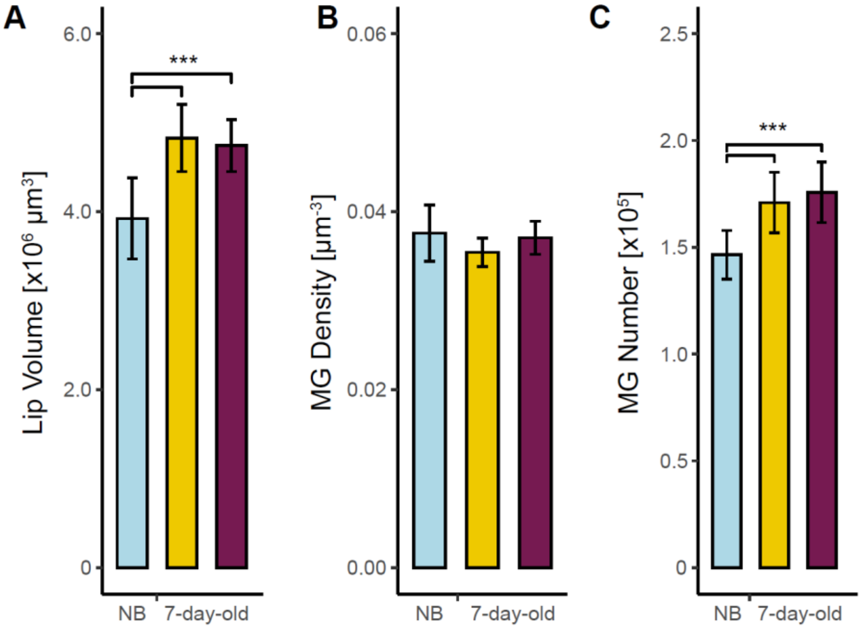


**Supplementary Figure S2.** Lip structure measurements in newborns (NB; blue) and 7-day-old control (yellow) or imidacloprid-treated (5 ppb; violet) bees. **(A)** The lip volume significantly increased during the first week of adulthood and was not affected by the imidacloprid treatment. **(B)** The density of MG in the lip did not differ between groups. **(C)** The number of MG in the lip significantly increased with age and was not affected by the imidacloprid treatment. Error bars represent the SEM; (***) $p < 0.0005$; $n = 7$-14 per group.

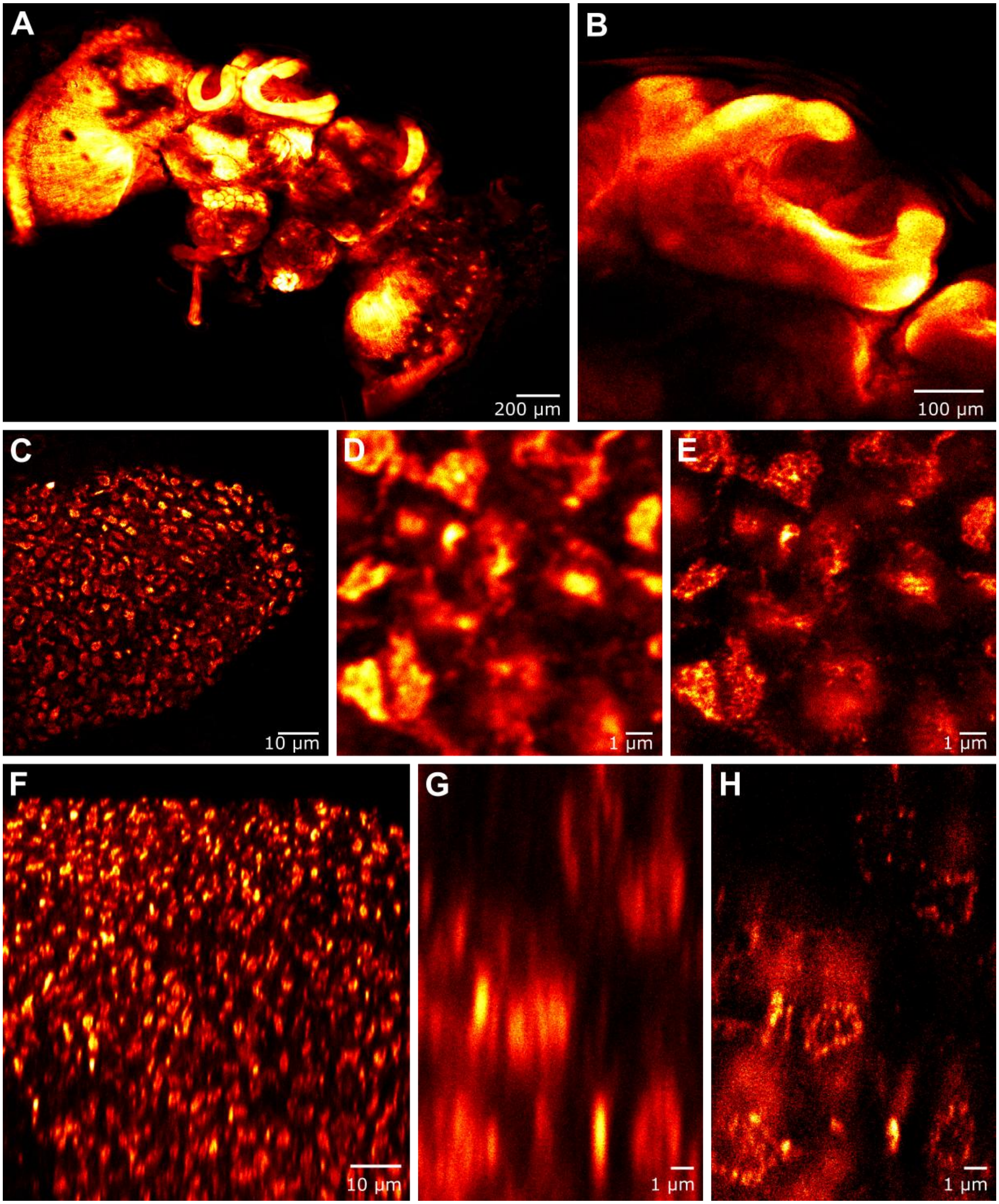


**Supplementary Figure S3. Optical sections of different honey bee brain regions following α-synapsin immunostaining of whole-mount brains.** (**A-E**) Horizontal cross-sections of the whole brain (**A**), the mushroom body neuropil (**B**), and the microglomeruli within the mushroom body lip (**C-D**), acquired with a confocal microscope. (**E**) STED image of the same region as (**D**). (**F-G**) Vertical cross-sections of the microglomeruli within a mushroom body lip subregion, acquired with a confocal microscope, (**H**) 3D-STED image of the same region as (**G**).

**Supplementary Video 1. Slice-through volume animation.** Image stacks of microglomeruli within a lip subregion were acquired with a confocal (left) and a STED (right) microscope. Images are from whole-mount brains immunostained for synapsin. The STED images were filtered using a Difference-of-Gaussians (DoG) filter. Scale bar is 1 µm.

**Supplementary Video 2: 3D rendering animation of a single microglomerulus.** The 3D volume of a microglomerulus was reconstructed using image stacks acquired in confocal (left) and in STED (right) mode. Images are from whole-mount brains immunostained for synapsin. The STED images were filtered using a DoG filter before the volume reconstruction. They resolve synapsin-positive units (SPUs) within the microgomerulus. Scale bar is 500 nm.
